# A synthetic RNA-mediated evolution system in yeast

**DOI:** 10.1101/2021.02.27.433199

**Authors:** Emil D. Jensen, Marcos Laloux, Beata J. Lehka, Lasse E. Pedersen, Tadas Jakočiūnas, Michael K. Jensen, Jay D. Keasling

**Affiliations:** Novo Nordisk Foundation Center for Biosustainability, Technical University of Denmark, Kgs. Lyngby, Denmark; Joint BioEnergy Institute, Emeryville, CA, USA; Biological Systems and Engineering Division, Lawrence Berkeley National Laboratory, Berkeley, CA, USA; Department of Chemical and Biomolecular Engineering & Department of Bioengineering, University of California, Berkeley, CA, USA; Center for Synthetic Biochemistry, Institute for Synthetic Biology, Shenzhen Institutes of Advanced Technologies, Shenzhen, China

## Abstract

Laboratory evolution is a powerful approach to search for genetic adaptations to new or improved phenotypes, yet either relies on labour-intensive human-guided iterative rounds of mutagenesis and selection, or prolonged adaptation regimes based on naturally evolving cell populations. Here we present CRISPR- and RNA-assisted *in vivo* directed evolution (CRAIDE) of genomic loci using evolving chimeric donor gRNAs continuously delivered from an error-prone T7 RNA polymerase, and directly introduced as RNA repair donors into genomic targets under either Cas9 or dCas9 guidance. We validate CRAIDE by evolving novel functional variants of an auxotrophic marker gene, and by conferring resistance to a toxic amino acid analogue in baker’s yeast *Saccharomyces cerevisiae* with a mutation rate >3,000-fold higher compared to spontaneous native rate, thus enabling the first demonstrations of *in vivo* delivery and information transfer from long evolving RNA donor templates into genomic context without the use of *in vitro* supplied and pre-programmed repair donors.

## Introduction

The ability to evolve biomolecules with tailor-made properties is inherently linked to mutagenesis, driving both natural and laboratory evolution. However, with the extreme high fidelity of genome replication, occurring with mutational frequencies in the order of one mutation per billion replicated DNA bases (i.e. 10^−9^ per base) (1), a multitude of directed evolution systems have been developed to increase both mutation rates and targeted mutation space (2, 3). While the vast majority of these systems rely on targeted mutagenesis of genomic loci using variant DNA donors designed and generated *in vitro* (4–7), a number of evolution systems have been developed to couple mutation and selection cycles *in vivo* in both bacteria (2, 8–11), yeast (12– 15), and mammalian cells (16). Such strategies circumvent the need for repeated cycles of human-guided design of mutational spectra, tedious hands-on genetic library construction, transformation, and selection, and have enabled targeted per-base substitution rates more than 10,000-fold higher than those of host genomes (e.g. 10^−5^ - 10^−4^ per base) (14, 17, 18).

Importantly, when developing systems for directed evolution *in vivo*, orthogonal mutagenesis and subsequent targeted delivery of mutant donors is of primary importance, in order to efficiently dereplicate sequence to function under selective conditions (19, 20). To address these considerations, creative bioprospecting and mixing of biological parts from diverse hosts have proven successful, including delivery of DNA mutant donors by heterologous faulty DNA polymerases and targeted base-editing using protein-fusions strategies (9–11, 13, 14, 16). Interestingly, various viral phylogenies store genetic information with low replicative fidelity (up to 10^−4^ per base per infection) in the form of RNA-encoded genomes (21), and viral-derived components have been a rich source for prospecting parts for synthetic directed evolution systems (8, 10, 22, 23). Moreover, RNA has been shown to serve as direct templates for DNA double-strand break (DSB) repair by homologous recombination *in vitro* and in yeast, and later also in bacteria and human cell lines (7, 24–27). Likewise, it has been demonstrated that RNA molecules synthesized *in vivo* can confer genome editing following induced DSBs (28, 29).

Based on this, RNA constitutes an interesting entry-point for development of directed evolution of DNA through RNA *in vivo*, yet this requires controlled delivery of diversified RNA donors to be established and means to target them to genomic loci of interest. Here we report the development of a synthetic *in vivo* directed evolution system for yeast using CRISPR/Cas9 or nuclease-deficient dCas9 (30–33) technology for RNA-programmed targeting of genomic loci with evolving chimeric donor gRNAs (cgRNAs) continuously delivered from an engineered low-fidelity T7 phage-derived RNA polymerase (T7RNAP). In this study, we first establish an inducible CRISPR/Cas9 system with cgRNA that allows for studying cgRNA-DNA repair in yeast. Next, we report the engineering and optimization of controlled and orthogonal delivery of multiple cgRNAs, and we demonstrate that the CRISPR- and RNA-assisted *in vivo* directed evolution (CRAIDE) system supports a mutation rate >3,000-fold higher than native replication fidelity at user-defined genomic loci, thus providing the first example of an RNA-based directed evolution system *in vivo*.

## Results

### Engineering orthogonal cgRNA delivery in yeast

In order to develop a targeted *in vivo* evolution system, we initially sought to combine elements of RNA-programmed genome targetability of CRISPR/Cas9, and error-prone RNA polymerase for expression of donor-coupled chimeric gRNAs (cgRNAs), serving as repair templates at targeted genomic loci (31, 32, 34). For choice of RNA polymerase, we selected bacteriophage T7RNAP, originally reported to produce mRNA transcripts, and more recently also functional gRNAs, in yeast (35, 36). Importantly, beyond orthogonal transcription relying on the high T7 promoter-specificity and synthesis of untranslated RNA in yeast by T7RNAP (37), transcriptional mutagenesis can be adjusted by evolved T7RNAPs with nucleotide substitution error rates up to 1.25 × 10^−3^ demonstrated *in vitro* and in *E. coli* (38), making T7RNAP of particular interest for *in vivo* evolution.

From this design, we first evaluated genome editing efficiency at the *ADE2* locus using wild-type T7RNAP in combination with *Streptococcus pyogenes* Cas9, and an *ADE2* gRNA (Fig. 1A, Suppl. Fig. S1). When co-transforming a 90-mer double-stranded DNA oligo (dsOligo) to knock-out *ADE2* we observed modest 2% genome editing efficiency, whereas leaving out dsOligo lowered efficiency to 0.04%, while no *ADE2* disruption was observed when both T7RNAP and dsOligo were omitted (Fig. 1A, Suppl. Fig. S1). To investigate *in vivo* delivery of RNA-mediated repair templates we next constructed chimeric donor gRNA (cgRNA) based on a 200 nucleotide 5′-primed extension of gRNA homologous to *ADE2* with PAM site and four PAM-proximal seed bases omitted to safeguard target site from repetitive cutting and frameshift-induced knock-out following cgRNA-templated DSB repair, respectively (Fig. 1B, Suppl. Fig. S1). We also tested a Cas9 variant reported to have improved genome editing efficiency (iCas9: Cas9^D147Y,P411T^)(39). Indeed, from co-transformations of T7RNAP and cgRNA together with either iCas9 or Cas9, we obtained 86% and 6% gene editing, respectively (Fig. 1B). However, in both cases, background gene editing efficiencies when T7RNAP was omitted reached 43% and 3%, indicating leaky cgRNA expression from the first-generation plasmid design (pEDJ350; Fig. 1B).

**Fig. 1.**
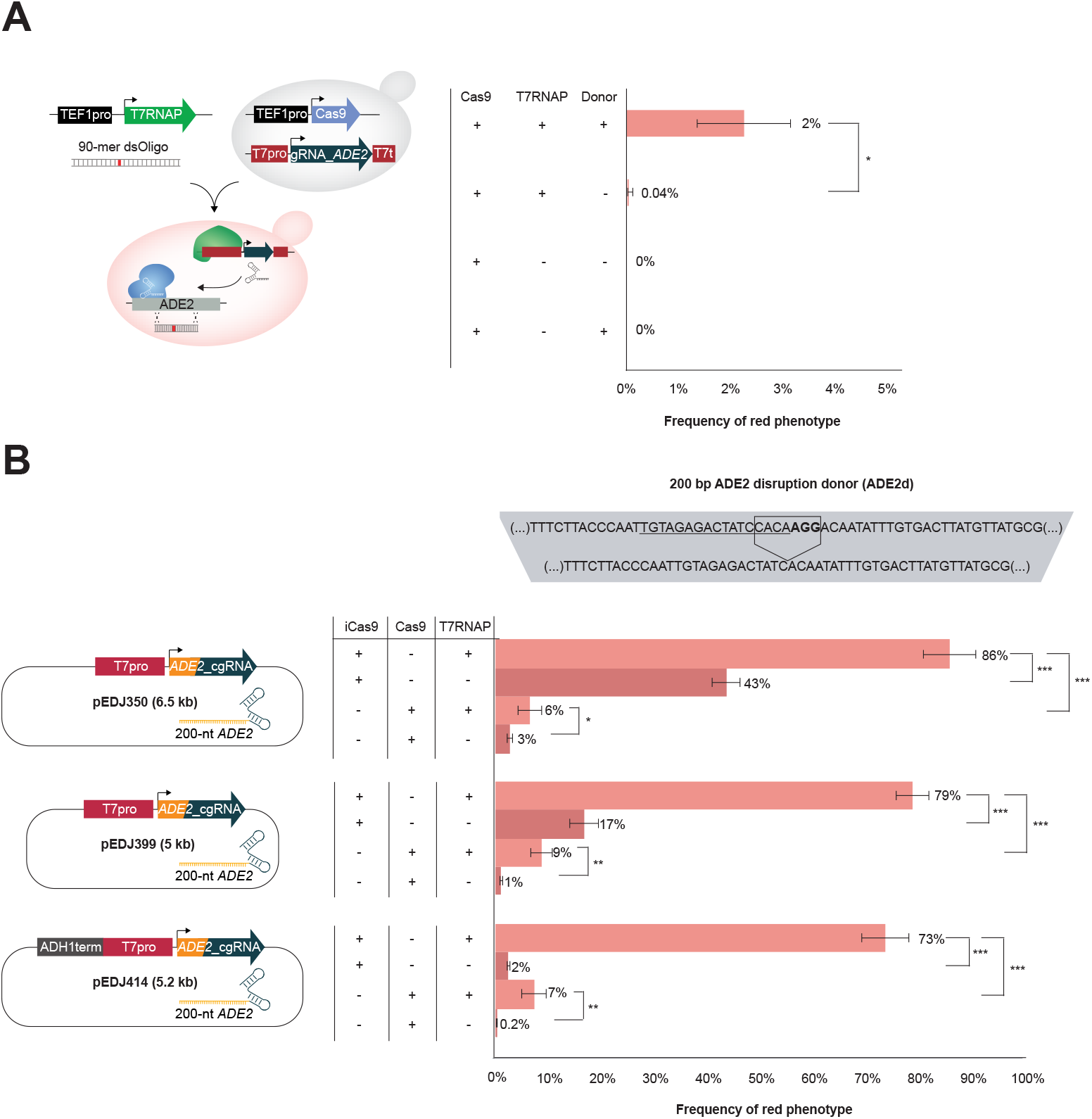
T7 RNA polymerase controls functional expression of gRNA and chimeric gRNAs (cgRNAs) in yeast. (**A**) Schematic illustration of experimental set-up, and frequencies of red colonies as a proxy for *ade2* knock-out and adenine deficiency. T7 RNA polymerase (T7RNAP) is indispensable for red colony formation in yeast cells expressing Cas9 under the control of the constitutive *TEF1* promoter (TEF1pro) and gRNA under the control of the T7 promoter (T7pro), when co-transformed with a linear double-stranded oligo (dsOligo) targeting disruption of *ADE2*. Mean frequencies of red colony formation ± s.d. from three (*n* = 3) biological replicate experiments. (**B**) Frequencies of red colonies in yeast cells expressing chimeric gRNA (cgRNA) targeting *ADE2* (*ADE2*_cgRNA). The cgRNA is based on a 200 nucleotide 5′-primed extension of gRNA homologous to *ADE2* with PAM site and four PAM-proximal seed bases deleted (disruption donor; ADE2d). Frequencies from yeast cells expressing either improved Cas9 (iCas9) or Cas9 in the presence or absence of T7RNAP are shown as mean ± s.d. from three (*n* = 3) biological replicate experiments.

As orthogonal and controlled delivery of evolving cgRNAs by T7RNAP is of paramount importance for practical applications, we mitigated high background gene editing by i) removing unannotated sequences in the cgRNA expression plasmid targeting *ADE2* (pEDJ399; Fig. 1B), and ii) introducing Pol II RNAP terminator from *ADH1* gene (ADH1t) upstream the T7 promoter on the pEDJ399 plasmid (pEDJ414; Fig. 1B). From these two approaches, background gene editing was lowered to 17% and 2% for iCas9, and to 1% and 0.2% for Cas9 (Fig. 1B), while at the same time maintaining T7RNAP-mediated gene editing efficiencies of 73-79% and 7-9% for iCas9 and Cas9, respectively. Moreover, inserting Pol III RNAP terminator SUP4 (SUP4t) upstream of the T7 promoter in pEDJ350 did not affect background expression, whereas inserting ADH1t in pEDJ350 reduced background expression similarly to the reduction observed in pEDJ414, indicating that the insulating properties of the ADH1t are indispensable for tight control of cgRNA expression (Suppl. Fig. S2).

In summary, we established an adjustable genome engineering system based on Cas9 variants and orthogonal delivery of functional cgRNA.

### Repair of plasmid DNA by cgRNA

To demonstrate that DSBs are repaired by T7RNAP-mediated delivery of cgRNA, and not by DNA-DNA homologous recombination between the cgRNA-expressing plasmid and the genomic target locus, we leveraged a previously established system to study transcript-mediated DSB repair (40). In this system, spliced antisense *HIS3* transcripts can serve as homologous templates to repair DSBs in the *HIS3* ORF interspersed by an artificial intron (*AI*), and subsequently allow for conditional expression of native *HIS3* transcripts read in sense orientation (Fig. 2A) (40, 41). In our modified system we initially fused cgRNA 3’-end of antisense *HIS3* (*HIS3_AI_cgRNA*) expressed under the control of the T7 promoter and introduced this plasmid into cells with T7RNAP and Cas9 expression induced by galactose. We used this design to test if CRISPR/Cas9-mediated DSB in the plasmid could be repaired by spliced *HIS3_AI_cgRNA* transcripts originating from the plasmid itself (*cis*). An early committed step for RNA-mediated repair of DSB is the formation of RNA-DNA duplexes, and RNase activity has been shown to inhibit RNA-DNA repair in eukaryotes (40, 42). For this reason we tested T7RNAP-mediated delivery of *HIS3_AI_cgRNA* in both wild-type cells and in cells deleted for RNase H1 (*RNH1*) and RNase H2 (*RNH201*) (40). Using replica-plate workflows we grew up wild-type and *rnh1 rnh201* cells with glucose, then replicated colonies onto galactose or glucose, and finally onto selective media without histidine to score colony-forming units following 3 days cultivation (Fig. 2A). When inducing expression of T7RNAP and Cas9 in *rnh1 rnh201* cells, 36% of the colonies turned histidine prototrophic (pEDJ367; +galactose), whereas only 0.1% of colonies from glucose control medium survived without supplemented histidine (Fig. 2B). Furthermore, from galactose-induction medium, the number of histidine prototrophic colonies drastically decreased to 0.2% and 3% from cells with deletions of either T7 promoter or cgRNA in the *HIS3_AI_cgRNA* expressing plasmid, respectively, and no colonies appeared without induction (Fig. 2B). Finally, we never detected any colonies on selective medium following induction of Cas9 and T7RNAP in wild-type cells, and neither did we observe any colonies from cells without T7RNAP (Fig. 2B).

**Fig. 2.**
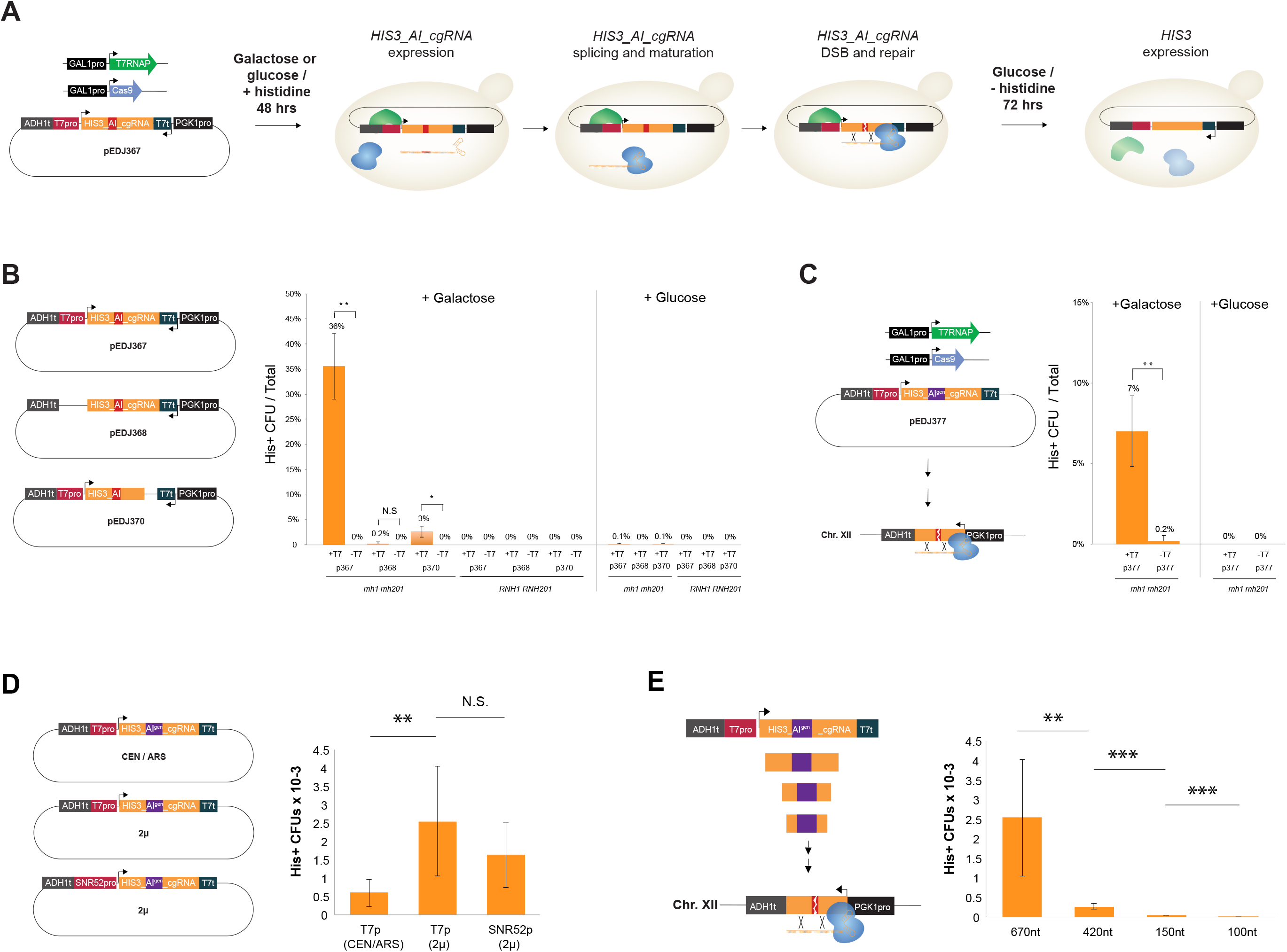
Cas9-mediated DNA double-strand breaks are repaired by RNA donors encoded in chimeric guide RNAs. (**A**) Schematic outline of the inducible replica-plating work-flow used for inferring repair of DNA double-strand breaks by donor RNA encoded in cgRNAs. (**B**) Dual-expression 2μ plasmid-based designs co-transformed into yeast together with Cas9 alone or with both T7RNAP and Cas9, show that inducible expression of T7RNAP enables efficient repair of Cas9-mediated DSB in the plasmid-encoded (*cis*) artificial intron (*AI*) positioned in the *HIS3* open reading frame when expressed in RNase-deficient (*rnh1 rnh201*) yeast. His^+^ colony forming units (CFUs) out of total colonies are shown. (**C**) Cas9-mediated DSB of *HIS3_AI*^*gen*^ in a single-copy genome-encoded (*trans*) *his3 AI*-disrupted reading frame (Sc71) can be repaired by donor RNA encoded in cgRNAs expressed by inducible T7RNAP in RNase-deficient (*rnh1 rnh201*) yeast. (**D**) cgRNA expression impacts cgRNA-DNA repair efficiency. A liquid assay was conducted with *rnh1 rnh201* strains with the cgRNA construct from pEDJ377 contained in centromeric (CEN/ARS) or 2μ plasmids and expressed from T7 promoter (T7pro) or SNR52 promoter (SNR52pro) as indicated. Genome-integrated *HIS3_AI*^*gen*^ was the target, and T7RNAP and Cas9 were inducibly expressed with galactose for 48 hrs prior to plating and His^+^ scoring. Colony-forming units (CFUs) were calculated relative to plating efficiency on non-selective media. (**E**) cgRNA-DNA repair with various donor sizes were investigated as in (D) by symmetric truncations of the cgRNA construct contained in pEDJ377 targeting genome-integrated *HIS3_AI*^*gen*^. For (B-E) frequencies of histidine prototrophic colonies and their error bars are shown as mean ± s.d. from three (*n* = 3) biological replicate experiments and significance determined from Student’s t-test, where * p<0.05, ** p<0.005, *** p<0.0005, and N.S. = not significant.

Taken together, these results highlight a tightly controlled cgRNA delivery system for transcript-mediated repair in RNase-deficient yeast.

### Repair of genomic DNA by cgRNA

Next, to enable a portable evolution system for delivery of candidate cgRNAs to target genomic loci, we determined if plasmid-based cgRNA expression could also support genome editing (*trans*). To enable this analysis, we changed 5 PAM-proximal bases (GAGTC) in the original cgRNA of the *HIS3_AI_cgRNA* plasmid into complementary bases (CTCGA, *HIS3_AI*^*gen*^), to specifically allow Cas9 to be guided to an integrated new synthetic *HIS3_AI* design matching seed sequence CTCGA only found in the genomic target locus (Fig. 2C). Repeating the workflow described above, induction of Cas9 and T7RNAP in *rnh1 rnh201* cells supported increased colony numbers (7%) under selective conditions, whereas control cells without T7RNAP expression only supported modest colony numbers (0.2%) under the same conditions, confirming that T7RNAP mediated expression of cgRNA directs Cas9 and templates DSB repair in genomic contexts (Fig. 2C).

To further test how cgRNA expression influences cgRNA-DNA repair, we induced cells in liquid dropout media and compared cgRNA expression from multicopy plasmids (2μ) and centromeric plasmids (CEN/ARS). Here, we found that using multicopy plasmids for cgRNA expression was >4-fold (*p* < 0.005) more efficient than expression from centromeric plasmids (Fig. 2D), whereas the use of more active native RNA polymerase III *SNR52* promoter(36) to drive cgRNA expression did not further improve cgRNA-DNA repair (Fig. 2D). This result indicates that cgRNA expression is not a limitation when expressed with T7RNAP from multicopy plasmids, and furthermore serves to illustrate that the cgRNA-DNA repair system can be scored based on simple liquid passaging.

Moreover, since homology size is paramount to efficient DNA-DNA repair(43), and has also been demonstrated to affect RNA-DNA repair(24), we next investigated cgRNA-DNA repair efficiencies of differently sized truncations of the cgRNA donor sequence compared to full-length donors (670 nt). Here, we found that longer homology regions (670 nt) were ∼86-fold more efficient for cgRNA-DNA repair compared to cgRNAs with short homology donors of 100 nt (Fig. 2E).

In summary, controllable plasmid-based cgRNA expression on plates or in liquid cultures can be designed to target the genome, where expression of long cgRNA donors from 2μ plasmids improves cgRNA-DNA repair efficiency.

### cgRNA-mediated directed evolution in vivo

Next, to investigate if DSB can be repaired by erroneous cgRNA donors, and thereby make way for establishment of RNA-mediated directed evolution in genomic contexts, we combined our established system for control of cgRNA delivery and Cas9-mediated targeting with the expression of a recently described error-prone T7RNAP double mutant (T7RNAP^F11L/T613A^) (22). T7RNAP^F11L/T613A^ was originally derived from a triple mutant with error-rates reported in *E. coli* studies to approximate 1.25 × 10^−3^ per transcribed base(38). However, though the triple-mutant did not express well in yeast, T7RNAP^F11L/T613A^ was observed to increase *ADE2* disruption over T7RNAP by 5-fold (Suppl. Fig. S3), and was therefore sought for evolving cgRNAs and genomic loci *in vivo*.

To test RNA-mediated directed evolution using T7RNAP^F11L/T613A^, we initially targeted resistance towards the toxic arginine analogue, L-Canavanine, as a proxy for genome evolution(36), by directing Cas9 to genomic *CAN1* using evolving 660-nt cgRNA donors (Fig. 3A). Following three days of directed evolution in liquid cultures, we scored mutation frequency based on canavanine-resistance observed in Cas9- and T7RNAP^F11L/T613A^-expressing cells either with or without the expression of *CAN1*_cgRNA. Here, we identified mutation frequencies of 1.8 × 10^−5^ +/- 1.3 × 10^−6^ and 3 × 10^−2^ +/- 6 × 10^−4^ for cells without and with cgRNA expressed, respectively, totalling 1,653-fold higher mutation frequencies in cgRNA-expressing populations compared to populations not expressing cgRNA (*p* = 1.14E-07) (Fig. 3B). Sequencing of the genomic *CAN1* locus identified K405N and S442Stop mutations in strains expressing T7RNAP^F11L/T613A^ with cgRNA and Cas9, with mutational spectrum spanning up to 107 bases from the DSB (Fig. 3C). None of the few colonies arising from strains lacking cgRNA had *CAN1* mutations within the donor region (Fig. 3C).

**Fig. 3.**
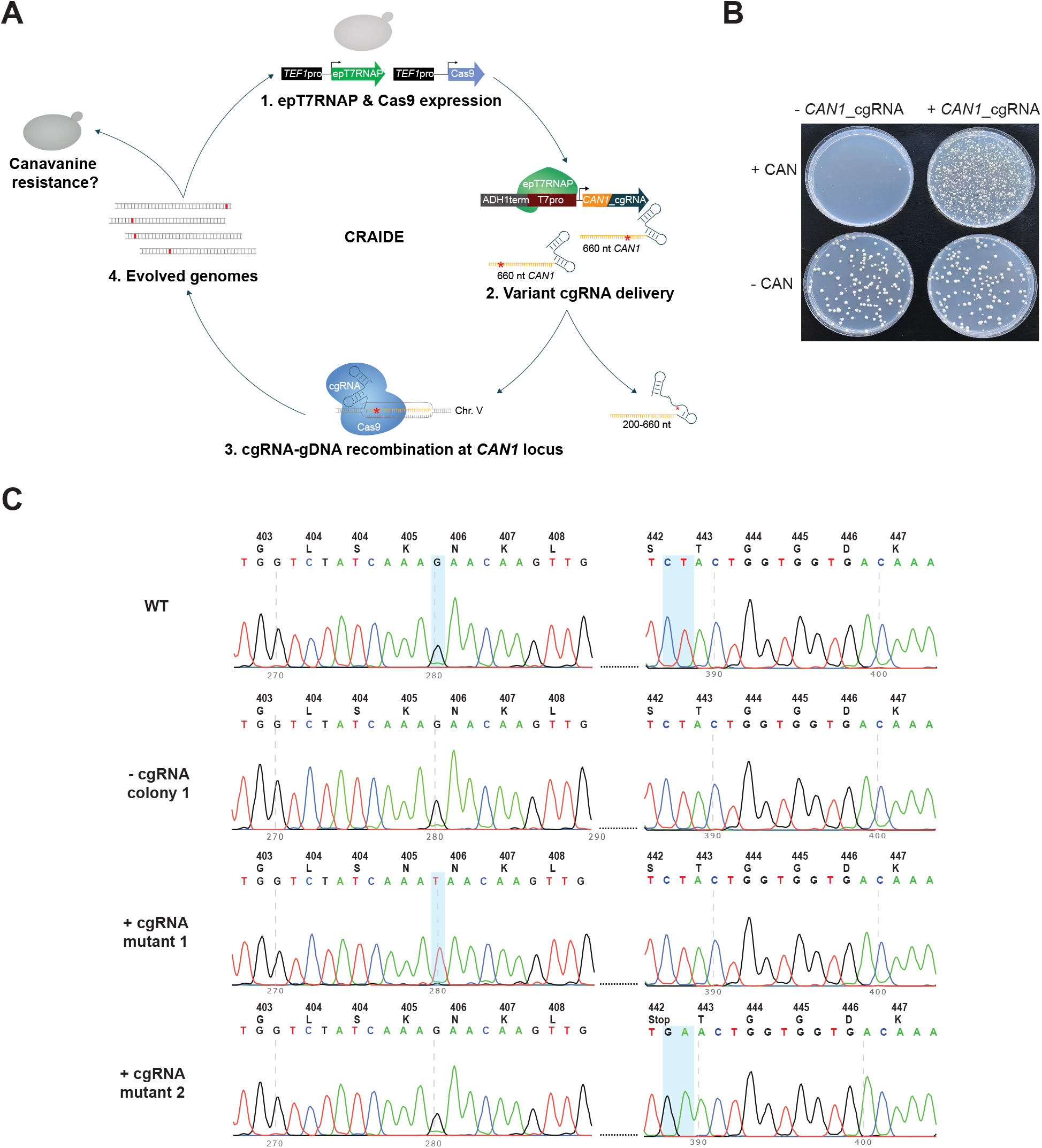
Orthogonal delivery of cgRNAs for targeted genome evolution *in vivo*. (**A**) Schematic outline of CRISPR-assisted programing of RNA-mediated *in vivo* directed evolution (CRAIDE). (**B**) Evolving resistance to L-Canavanine by error-prone T7RNAP^F11L/T613A^-mediated transcription of cgRNAs encoding a 660-nt donor for positive selection of *CAN1* disruption. Representative colony numbers on selective (+CAN) and non-selective (-CAN) plates. Following 3 days of evolution in liquid cultures, 50 μl of control (-cgRNA) and CRAIDE (+cgRNA) cultures were plated onto selective and non-selective plates, and mutation frequency scored based on numbers of L-Canavanine-resistant colonies on three (*n* = 3) biological replicates. For each plate 50 μl of saturated liquid culture was plated. For -CAN plates, the 50 μl was diluted 500x before plating. (**C**) Sanger-based sequencing of genomic *CAN1 locus* in WT cells and two canavanine-resistant (*can1*_mut.1 and *can1*_mut. 2) colonies from *CAN1*-cgRNA expressing CRAIDE cultures.

Encouraged by these results, and by the fact that mutagenesis associated with nuclease-deficient Cas9 (dCas9)(44) has been observed previously(45–47), we next sought to test if dCas9 could facilitate RNA-DNA editing without Cas9-induced DSB using the *CAN1* RNA-DNA repair screen, and 10X higher concentrations of L-Canavanine compared to Fig. 3 to diminish residual growth (Suppl. Fig. S4A and S4B). Here, Cas9 performed ∼3.5-fold better than dCas9 (*p* = 0.017) with resistant colonies appearing at a frequency of 2.2 × 10^−5^ and 6.3 × 10^−6^ in viable cells, respectively, after induction (Suppl. Fig. S4C), while dCas9 sustained higher cell densities (*p* = 0.031). By contrast, strains with no cgRNA expression appeared 229-fold less frequently on selective plates compared to when both Cas9 and cgRNA were expressed (9.6 × 10^−8^; *p* = 0.0045 and *p* = 0.011 for Cas9 and dCas9, respectively; Suppl. Fig. S4C). These results provide a first demonstration of using dCas9 for cgRNA-DNA editing.

Finally, to fully demonstrate the applicability of directed evolution with cgRNA-DNA repair, we tested CRAIDE for gain-of-function mutagenesis in a genomic locus. For this purpose, we targeted a genome integrated design (HIS3_23Δ29-XII-5) lacking 29 bases of the *HIS3* open reading frame, and hence rendering cells unable to grow without histidine supplementation. Here, galactose inducible T7RNAP^F11L/T613A^ and Cas9 were expressed together with *cgRNA_HIS3_stop* containing a STOP codon at *HIS3* position K71 (pEDJ508), which is surrounded by the 29 bp deletion in the genomic design to rule out the possibility of NHEJ repair in surviving mutants. More specifically, the *cgRNA_HIS3_stop* was engineered to contain a STOP codon (A211T; <u>A</u>AG-><u>T</u>AG) three bases upstream from the artificial intron and ten bases from the Cas9-generated DSB in HIS3_23Δ29-XII-5 (Fig. 4A). Hence, by design, only induced cells successfully repaired with a cgRNA which had the encoded STOP codon evolved into a permissive mutation would be able to sustain growth under selection (w/o histidine supplementation). Repeating the liquid passaging set-up as previously adopted (see Fig. 2D-E) we induced seventeen replicate cultures each transformed with the plasmid expressing *cgRNA_HIS3_stop* (pEDJ508), along six replicate control cultures transformed with empty no-cgRNA vector (pEDJ400), for 48 hrs under non-selective conditions (galactose, with histidine; Fig. 4B). Next, cultures were plated on histidine dropout media to score the mutation frequency, and for a subset also propagated in liquid non-inducing selective conditions (glucose, w/o histidine) for three days to score growth. Indeed, while cultures carrying plasmid pEDJ508 grew to saturation, replicate cultures carrying pEDJ400 did not grow (Fig. 4C), and neither did we observe any colonies on selective plates from cultures without cgRNA expression (Supplementary Table S1). Moreover, from amplicon sequencing of the repaired target site (i.e. HIS3_23Δ29-XII-5) of saturated cultures with pEDJ508, we found various mutations that abolished the STOP codon. Here, the replicate cultures carried one to three mutations containing either T>G and C>A, translating into STOP>E and H>N, CTA>TGT translating into STOP>V, or one distinct mutation A>C leading to STOP>S (Fig. 4D). To further substantiate the ability of the plasmid-based CRAIDE system to selectively target genomic loci of interest, we also sequenced plasmid pools (pEDJ508) from replicate cultures. Here, the pre-engineered STOP codon in *cgRNA_HIS3_stop* was observed in all of the cultures (Suppl. Fig. S5B). Importantly, reintroduction of identified STOP codon mutations from the repaired HIS3_23Δ29-XII-5 genomic locus into clean genetic background strains verified histidine prototrophy in all cases (Suppl. Fig. S5C). Thus, from this parallelized directed evolution study, CRAIDE conferred both transitions, transversions, and combinations thereof.

**Fig. 4.**
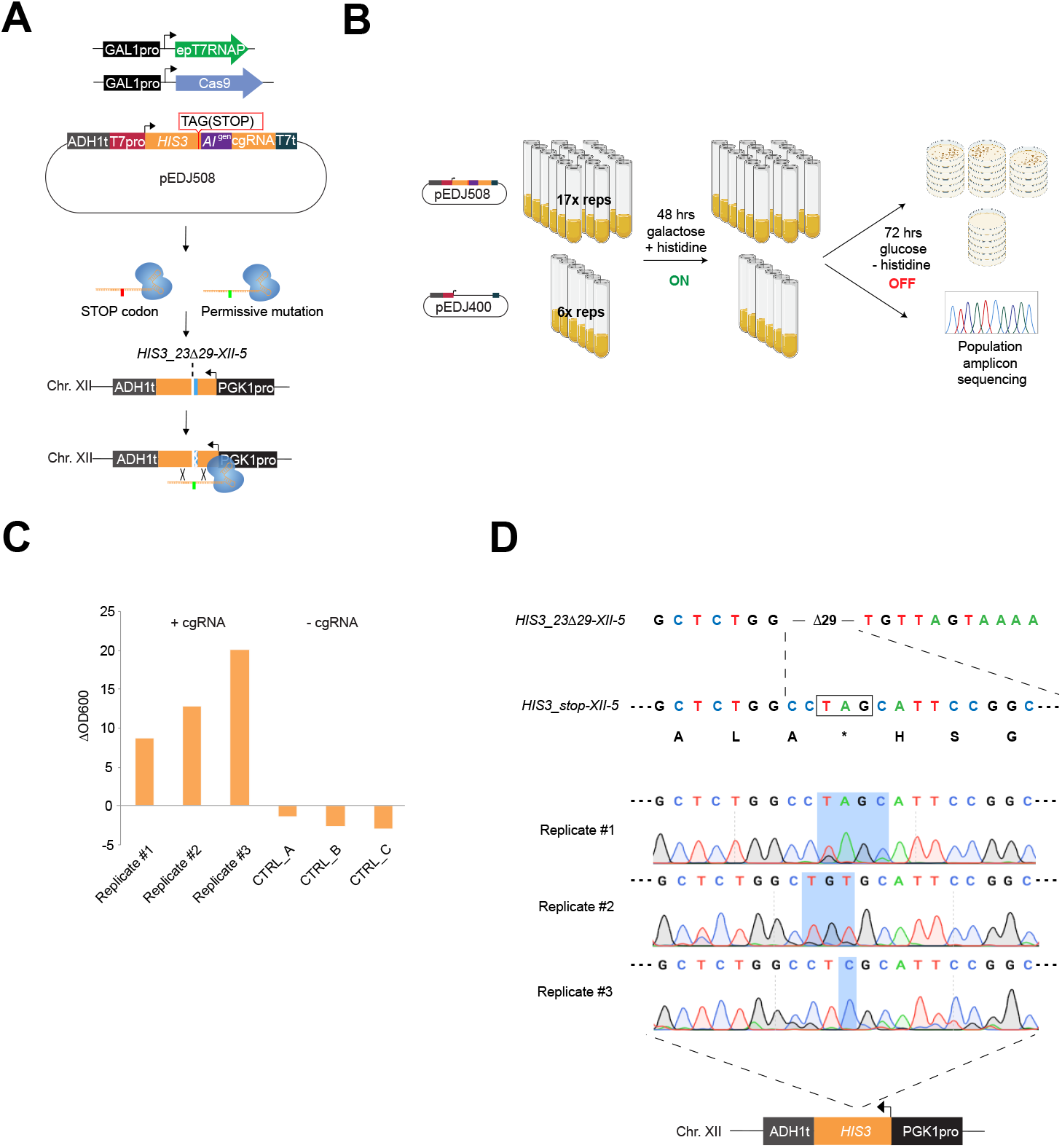
Directed evolution by cgRNA-DNA repair displays targeted transitions and transversion. **(A)** Plasmid-based galactose-inducible Cas9 and T7RNAP^F11L/T613A^ (epT7RNAP) were expressed with plasmid pEDJ508 (described in the main text). On system induction for 48 hrs in synthetic complete dropout media with galactose, *cgRNA_HIS3_stop* expressed from pEDJ508 directs Cas9 to the genome-integrated single copy HIS3_23Δ29-XII-5 cassette to induce DNA double-strand break and template DNA repair. The cgRNA may contain the engineered STOP codon sequence (red), or evolved permissive mutations (green) introduced with epT7RNAP. **(B)** Biological replicates were induced (ON) with galactose dropout media for 48 hrs in 2 ml volumes. 300 μl were plated for each replicate on five plates containing histidine dropout media with glucose to stop evolution (OFF). (**C**) Remaining cells were transferred into liquid histidine dropout media with glucose (OFF) and incubated for 72 hrs to determine growth as indicated by change in OD600 (ΔOD600) during the 72 hrs cultivation in triplicate cultures expressing either pEDJ508 (+cgRNA) or pEDJ400 (-cgRNA). (**D**) Sequencing results from amplicon sequencing of 500 μl of saturated liquid cultures (population level) expressing a repaired HIS3_23Δ29-XII-5 allele (as indicated schematically at the bottom). Corresponding amino acids are shown below *HIS3_stop-XII-5*, and TAG (STOP) is boxed and indicated by an asterisk (*). Chromatograms are given for biological replicates #1-3, where blue shading spans the range of mutated bases. Colony scores and OD600 values are presented in Supplementary Table S1.

Finally, based on colony numbers from growth under selective conditions (see Suppl. Table S1), the initial mutation rate was estimated to be 9.77 × 10^−6^ per cell per generation, and the per-base mutation rate was determined to be 3.26 × 10^−6^ by adjusting for the number of bases (3) that can give rise to a permissive codon after repair of HIS3_23Δ29-XII-5 (for detailed explanation see Methods section *Estimation of mutation frequency and rate*).

Taken together, from the genotyping of gain-of-function mutants, any base (A, T, C, or G) can be introduced into the cgRNA during transcription and further transferred into a targeted genomic sequence, thus establishing inducible directed evolution *in vivo* based on RNA-mediated genome editing, with a mutation rate of 3.26 × 10^−6^ per base, being >3,000-fold higher than native background mutation frequency (1). Importantly, no mutants appeared on selective plates or in liquid cultures without cgRNA expressed (Fig. 4C and Suppl. Table S1).

## Discussion

This study demonstrates RNA-mediated and CRISPR-guided *in vivo* editing and mutagenesis at targeted genomic loci. To enable this, we first optimized orthogonal control of cgRNA expression using T7RNAP and insulated T7 promoters, and next demonstrated cgRNA-DNA repair on targeted genomic DSBs generated with Cas9. Extending from these results, by using an error-prone variant of T7RNAP for *in vivo* delivery of cgRNAs with random mutations, we enabled the first demonstration of directed evolution based on long evolving RNA donor templates into genomic contexts using both Cas9 and dCas9, and without the use of *in vitro* supplied and pre-programmed repair donors, as routinely adopted in directed evolution systems (5–7, 48).

However, engineering *in vitro* and *in vivo* directed evolution systems has experienced a lot of attention since their first demonstrations landmarked by Wright & Joyce, and Esvelt *et al*., respectively (8, 23). For this reason, pros and cons should be addressed when developing novel directed evolution techniques. Here, compared to other *in vivo* directed evolution systems in yeast (14, 15, 17), limitations of the current version of CRAIDE exist and need consideration and further improvement for the system to be applicable for efficient *in vivo* directed evolution across multiple species. Indeed, with a mutation rate in the order of 3.26 × 10^−6^ per base, RNA-mediated repair of genomic contexts using variant RNA donors as demonstrated in this study is still 2-3 orders of magnitude less efficient compared to state-of-the-art *in vivo* directed evolution methods for bacteria, yeast, and mammalian cells, like OrthoRep, ICE and TRACE (11, 14, 16, 17, 49). Furthermore, even though RNA-mediated DNA repair has previously been reported in wild-type yeast (29, 40), in its current version, CRAIDE requires disruption of host RNases for successful RNA-mediated repair of genomic DSBs. Such genetic prerequisites restrict the immediate portability of the system to genetically tractable hosts for which RNase H disruption does not confer lethality (50). However, for such cases, one mitigation strategy could involve conditional mutants to relieve potential lethality or long-term genotoxicity. Likewise, whereas *S. cerevisiae* has a highly proficient homologous recombination machinery for DNA repair, other eukaryotes, including mammals, are biased towards NHEJ to repair genomic DSBs (51) and may undermine, or at least limit, cgRNA-DNA repair efficiency, which is an essential requirement for CRAIDE to function. However, as reported for mammalian cells, designing physical proximity between targeted genomic loci and gRNA-appended donors can limit such false-positive events (7). To avoid generation of indel mixtures from using Cas9 for genome editing (6), our successful demonstration of CRAIDE for genome editing using dCas9 should be of relevance for *in vivo* directed evolution in hosts with NHEJ-bias for DSB repair. Also, homologous recombination can be further prompted to facilitate CRAIDE in new hosts by directly fusing HDR enhancing proteins to Cas9 (52, 53). Moreover, target genes can be engineered prior to CRAIDE to completely avoid screening for mutants that result from NHEJ, by removal of bases adjacent to PAM, which are then subsequently re-introduced by cgRNA-DNA repair as was performed in this study. As more findings on mechanisms governing RNA-DNA repair emerge, new strategies, such as fusing DNA polymerase ζ or other polymerases possibly involved in RNA-DNA repair (54), are relevant to pursue.

Acknowledging these limitations and considerations, CRAIDE is still a complementary tool expanding the scope of existing *in vivo* directed evolution systems (14, 15, 17), and to the best of our knowledge the first to directly utilize erroneous RNA-templated DNA repair. Specifically, CRAIDE constitutes a versatile *in vivo* directed evolution system with tunability in terms of editing efficiency (e.g. cgRNA expression, length of donor), flexibility in terms of genomic target loci (Cas9-directed genic and intergenic regions), and mutational landscape determined by T7RNAP fidelity (any base, transversions and transitions), with a >100 bp editing window.

Lastly, it should be mentioned that during the preparation of this study, three complementary DNA-templated genome editing technologies were reported; prime editor, TRACE and T7-DIVA (7, 11, 16). Here, prime editor demonstrated RNA-mediated genome engineering using *in vitro*-edited donor-amended gRNAs (prime editing gRNAs) (7), while TRACE and T7-DIVA demonstrated that T7RNAP fused to base editors could be applied for continuous *in vivo* mutagenesis of target genes controlled by genomically integrated T7 promoters (11, 16). Individually, these new technologies enable >10^−4^ mutations per base in engineered T7pro-driven open reading frames sized up to 2 kb, and nuclease-deficient integration of mutant bases in a prime editor window of approximately 30 bases (7, 11, 16). In the future, we envision that the *in vivo* variant donor delivery and editing window size of CRAIDE together with the high editing efficiencies of these technologies could present appealing mergers for development of efficient *in vivo* continuous evolution in broad genomic contexts, as well as providing a tool for more foundational basic research on RNA-mediated evolution.

## Methods

### Molecular cloning

Oligonucleotides, gene block fragments, and double-stranded 90-mers were purchased from Integrated DNA Technologies (IDT). All primers, dsOligos, gRNAs, and plasmids constructed for this study are listed in Suppl. Table S2 and S3. Fragments for USER cloning were amplified with Phusion U Hot Start PCR Master Mix from ThermoFisher Scientific (catalogue #F533S), and assembly was done with USER enzyme (55) into *Sfa*AI/Nb.*Bsm*I-treated vectors as described previously(56). gRNA expression cassettes contained overhangs for cloning with universal USER-overhang primers previously described (57). Plasmid pEDJ8 expressing gRNAs for knock-out (KO) of *RNH1* and *RNH201* was constructed by assembling pre-synthesized gene blocks gEDJ1 and gEDJ2 into p0054 (58). Plasmid pEDJ332 was made by replacing the existing gRNA seed sequence in plasmid pCfB3050(59) to target *HIS3* (CAAGTGATTAACGTCCACAC). Cas9 was cloned from p414-Tef1p-Cas9-CYC1t (Addgene #43802) into vector pRS415U(60) with promoters from *GAL1* or *TEF1* to assemble pEDJ333 and pEDJ391, respectively. Cas9 nuclease mutants (pEDJ403-405 and pEDJ423) were made by site-directed mutagenesis. Wild-type T7RNAP was amplified from *E. coli* (DE3) with primers containing a truncated SV40 nuclear localization signal (NLS), and assembled with promoters from *TEF1* or *GAL1* into vector p0057 (58) to make pEDJ344 and pEDJ356, respectively. TEF1pro-T7RNAP was fused to GFP (amplified from gEDJ_GFP) in p0054 (pEDJ334) for expression analysis, and T7RNAP point mutants (pEDJ338-340, pEDJ342-343, pEDJ346, and pEDJ389) were generated by site-directed mutagenesis. epT7RNAP from pEDJ389 was then assembled with the *GAL1* promoter into p0057 to make pMLB10. Minimal vectors pEDJ400 and pEDJ437 were made by assembling auxotrophic markers (*URA3* or *HIS3*, respectively), ampicillin resistance gene (AmpR), origin of replication for yeast (2 micron) and bacteria (pUC), and USER cloning site. *ADE2* disruption cassette, including the T7 promoter (gEDJ3), and gRNA scaffold fused to the T7 termination signal (tZ)(61) (gEDJ4) were synthesized as gene blocks and assembled into p0054 (pEDJ350) or pEDJ400 (pEDJ399). pEDJ414 was made similarly by including ADH1t in the assembly. pEDJ372 was made by removing the *ADE2* disruption cassette from pEDJ350, and ADH1t was inserted into the *Eco*RI-*Sal*I restriction sites upstream from the expression cassette in pEDJ350 to make plasmid ADH1t_pEDJ350. SUP4t was inserted into pEDJ350 in both orientations by inverse PCR to make plasmids sup4tF_pEDJ350 and sup4R_pEDJ350. Antisense *HIS3_AI* expressed from the T7 promoter (gEDJ5) was assembled with ADH1t, gRNA:tZ, and PGK1 promoter to constitute plasmid pEDJ367. T7 promoter was excluded (pEDJ368) by inverse PCR, or gRNA scaffold was omitted from assembly (pEDJ370). pCfB2909(59) contains an integration cassette for EasyClone site XII-5(56) in yeast chromosomal DNA. Modified *HIS3_AI*^*gen*^ expressed from T7 promoter was synthesized as a gene block (gEDJ6) and assembled with ADH1t and *PGK1* promoter into pCfB2909 to make pEDJ375. Vector pEDJ377 was constructed by removing the *PGK1* promoter from pEDJ367 and modifying the seed sequence from TGTTAGTAAAAATTC<u>GAGCT</u> to TGTTAGTAAAAATTC<u>CTCGA</u> (change is underlined) by inverse PCR to match the artificial intron sequence residing in integrated pEDJ375. pEDJ465 was made by cloning an amplified fragment, comprising 660 nt of the wild-type *CAN1* sequence, with a ADH1t:T7pro fragment and gRNA:tZ into vector pEDJ400 to express cgRNAs against *CAN1*. Plasmid pEDJ423 was made by inverse PCR with primers F-Cas9(D10A) + F-Cas9(D10A) on pEDJ391, and then with primers F-Cas9(H840A) + F-Cas9(H840A) on the resulting plasmid. pMLB15 was made by ligation after inverse PCR on pEDJ375 with primers EDJ610+EDJ611, and plasmid pEDJ506 was made by ligation after inverse PCR on pMLB15 with primers EDJ610+EDJ654. pEDJ508 was made by ligation after PCR with EDJ657+EDJ658 on pEDJ377, and the design was then transferred to pEDJ400 with primers F-ADH1t-T7p + R-tZ. pMLB2 to pMLB4 were made by assembly into p0054 of 100bp/150bp/440bp_HIS3_AI(CTCGA) amplified from pEDJ377 with primers MLB1 to MLB6, together with fragments ADH1t:T7pro and gRNA:tZ as before. pMLB7 and pMLB8 were made by assembly into p0054 of HIS3_AI(CTCGA) amplified from pEDJ377 with primers MLB23+MLB24, gRNA:tZ or SUP4t, respectively, and ADH1t:T7pro or ADH1t:SNR52pro, respectively. pEDJ509 to pEDJ513 were made by ligation after inverse PCR of pEDJ437 with primers EDJ661 + EDJ662-666, respectively.

All strains, plasmids, oligos, gRNAs, and gene blocks are listed in Supplementary Tables 2-6.

### Baseline strain construction

Strain CEN.PK2-1C was chemically transformed with Cas9 (pEDJ391) to make baseline strain Sc35. Strain Sc36 was made by genetic deletion of *RNH1* and *RNH201* in Sc35 with 1 nmol double-stranded 90-mer oligos and 1 μg og RNA plasmid pEDJ8. Native *HIS3* was completely removed from Sc36 (Sc40) and Sc35 (Sc41) with 1 nmol of double-stranded 90-mer oligo and 1 μg gRNA plasmid pEDJ332. Strain Sc36 and Sc40 transformed with pEDJ391 were saved as Sc42 and Sc43, respectively. Sc43 was transformed with 3 μg *Not*I-linearized pEDJ375 or pEDJ506 (HIS3_23Δ29-XII-5) with 1 μg gRNA plasmid pCfB3050 to make Sc71 and Sc138, respectively.

### ADE2 disruption analysis

All media contains 2% glucose. CEN.PK2-1C was co-transformed with Cas9 (pEDJ391) or iCas9 (pCT; Addgene #60620) and T7RNAP (pEDJ344) (Sc104 and Sc106, respectively), or an empty vector w/o T7RNAP (p0057) (Sc103 and Sc105, respectively). Sc103-106 were incubated O/N in SC-Leu-Trp, then diluted 10X and incubated for 4 hrs at 30 °C with shaking prior to chemical transformation with relevant gRNA or cgRNA expression vectors. Sc103-104 were co-transformed with pEDJ372 and double-stranded 90-mer oligos with flanking homology to the *ADE2* break site. Transformed cells were resuspended in 100 μl mQ water and transferred into 3 ml liquid SC-LWU in a 15 ml culture tube and incubation at 30 °C for 72 hrs with shaking. Dilution series were plated after 72 hrs on SC-LWU, and red/white ratios for ∼500 colonies per plate were scored after 3 days of incubation at 30 °C. Chimeric red-white striped colonies (<5% per plate) were not considered.

### Replica-plating analyses of HIS3 repair

Incubations were performed at 30 °C. Sc40 and Sc41 were transformed with galactose-inducible T7RNAP (pEDJ356), Cas9 (pEDJ333), and pEDJ367, pEDJ368, or pEDJ370 (Sc107-109 and Sc113-115 for Sc40 and Sc41, respectively). Control strains carried p0057 instead of pEDJ356 (Sc110-112 and Sc116-118 for Sc40 and Sc41, respectively). Transformed cells were plated on SC-LWU with 2% glucose. Isolated colonies were inoculated in 1 ml of SC-LWU with 2% glucose and grown O/N with shaking. Dilution series were plated on the same media and incubated for 2 days prior to replica-plating. Replica-plating was done on SC-His (to adjust for rare spontaneous conversions) and on SC-LWU with 2% glucose or 2% galactose. Plates were incubated for 48 hrs before transferring to SC-His, followed by 3 days incubation. Sc71 was co-transformed with pEDJ377, galactose inducible Cas9 (pEDJ333), and T7RNAP (pEDJ356) (Sc119) or p0057 (Sc120). The same replica-plate workflow as used above for scoring His^+^ units was followed.

### Liquid induction analyses of HIS3 repair

Incubations were performed at 30 °C at 250 rpm. Strain Sc71 was transformed with pEDJ333, pEDJ356, and pEDJ377, pMLB2, pMLB3, or pMLB4 (Sc139 to Sc142, respectively) for donor-size analysis, and pEDJ333, pEDJ356, and pEDJ377, pMLB7, or pMLB8 (Sc143 to Sc145, respectively) for cgRNA expression analysis. 1 ml of each saturated culture was pelleted, washed once in 500 μl sterile mQ water, and resuspended in 200 μl sterile mQ water that was transferred to 3 ml of SC-LWU with 2% galactose (OD_600_ ∼2.0) for 48 hrs induction. Cultures were adjusted to OD_600_ = 2.0, and 3 × 1 ml were plated on SC-His, and serial dilutions plated on SC-LWU with 2% glucose.

### Gain-of-function analyses of HIS3_23Δ29-XII-5 repair

Incubations were performed at 30 °C at 250 rpm. Sc146 and Sc147 were made by transforming Sc138 with pMLB10, pEDJ333, and pEDJ508 or pEDJ400 (ctrl), respectively. Isolated colonies were inoculated for each strain in 5 ml of SC-LWU with 2% glucose for growth O/N. O/N cultures were washed once in sterile mQ water and and adjusted to OD_600_ ∼2.0 in 2 ml SC-LWU with 2% galactose and incubated for 48 hrs. Final OD_600_ was determined for Sc146 and Sc147, before plating 300 _μ_l on 5 plates of SC-His and dilutions on SC-LWU both with 2% glucose for each replicate culture. The remaining culture for three replicates was added 5 ml of SC-His with 2% glucose, and ΔOD_600_ was determined after 72 hrs of incubation. 500 μl of saturated cultures was harvested by boiling with 400 mM LiAce and 1% SDS for 10 min followed by ethanol precipitation and was finally resuspended in 100 μl mQ water. Amplicons were obtained by PCR with 2xOneTag master mix (ThermoFisher Scientific #K01s71) and primers MLB26+EDJ315 (genome) and EDJ360+EDJ353 (plasmid) and sequenced with the forward primer for each reaction.

### CAN1 survival assay

All media contained 2% glucose. Sc36 was transformed with epT7RNAP (pEDJ389) and Cas9 (pEDJ391) or dCas9 (pEDJ423) to make Sc127 and Sc134. Sc127 and Sc134 were transformed with pEDJ400 or pEDJ465 to give Sc128 and Sc129 for Sc127, respectively, and Sc135 and Sc136 for Sc134, respectively. Auxotrophies in Sc36 were closed by co-transformation of pRS415 (*LEU2*), p0057 (*TRP1*), and pEDJ400 (*URA3*). Biological replicates were inoculated in 5 ml SC-LWU and grown cultivated for 72 hrs in 15 ml culture tubes. Saturated cultures were plated on Delft supplemented with 20 mg L-Histidine (Delft+) and Delft+ supplemented with 60 μg/ml (in Fig. 3) or 600 μg/ml (in Suppl. Fig. S4) L-Canavanine to apply selection for *can1* mutants. 50 μl from three biological replicates +/-cgRNA were pelleted and supernatant discarded. Pellets were resuspended in water and plated, and the ratio of viable cells between strains expressing +/- cgRNA was determined after 3 days of incubation at 30 °C. Resulting genotypes from single colonies were determined by Sanger sequencing. Colony PCR was done with 2xOneTag master mix and primers F-*CAN1*-Sanger and R-*CAN1*-Sanger to amplify endogenous *CAN1* for sequencing analysis.

### Estimation of mutation frequencies and rate

All data and calculations are presented in Supplementary Table S1. Mutational frequencies were obtained by scoring the number of resulting colonies on selective media following evolution. The average number of mutants was then divided by the number of viable cells per plated volume for each culture (300 μl for gain-of-function, 500 μl for loss-of-function). Viable cells per volume were estimated from dilution series on non-selective media, and for gain-of-function analysis, the number of generations during 48 hrs system induction was determined from ΔOD_600_. Gain-of-function mutation frequencies were divided by the number of underwent generations, and then by three to adjust for the space that allows for permissive mutations in the STOP codon (TAG) in *HIS3_23Δ29-XII-5* after repair with mutant *cgRNA_HIS3_stop* (pEDJ508).

Combined this estimates a mutation rate of 3.26 × 10^−6^ per viable cell per generation per base. By comparison, commonly used(17, 22) online tools such as *bz-*rates (62) and rSalvador (63) estimated comparable mutation rates of 2.80 × 10^−6^ and 2.22 × 10^−6^, respectively. Yet, as *bz*-rates and rSalvador assume neglectable low starting ODs, while CRAIDE requires starting OD600 ∼2.0, we consider the mutation rate of 3.26 × 10^−6^ per viable cell per generation per base most accurate.

### Flow cytometry analysis

Strains Sc121-126 and p0054 were diluted 1:10 from O/N cultures into fresh 500 μl SC-Ura and incubated at 30 °C with shaking for 24 hrs prior to analysis. Cultures were diluted 1:5 in 150 μl with Phosphate Buffer Saline (PBS) from Life Technologies immediately before analysis by flow cytometry on the BD LSR Fortessa X-20 (BD Biosciences). Blue laser at 488 nm was used to analyse 10,000 single cells for each population, and FlowJo software (TreeStar Inc.) was used to process data and to calculate arithmetic mean fluorescence intensity values.

### Statistical analysis

Significance was determined by two-tailed Student’s t-test using at least three biological or technical replicates.

### Media

1 L of mineral media (Delft) with 2% glucose (64) contained 75 ml (NH_4_)_2_SO_4_ (100 g/L), 120 ml KH_2_PO_4_ (120 g/L), 10 ml MgSO_4_, 7H_2_O (50 g/L), 2 ml trace metals, 1 ml vitamins, and 20 g glucose. 1 L of trace metals contain 4.5 g CaCl_2_·2H_2_O, 4.5 g ZnSO_4_·7H_2_O, 3 g FeSO_4_·7H_2_O, 1 g H_3_BO_3_, 1 g MnCl_2_·4H_2_O, 0.4 g Na_2_MoO_4_·2H_2_O, 0.3 g CoCl_2_·6H_2_O, 0.1 g CuSO_4_·5H_2_O, 0.1 g KI, and 15 g EDTA. 1 L of vitamins contain 50 mg biotin, 200 mg p-aminobenzoic acid, 1 g nicotinic acid, 1 g Ca-pantotenate, 1 g pyridoxine HCl, 1 g thiamine HCl, and 25 g myo-Inositol. Synthetic complete dropout media were bought from Sigma-Aldrich.

## Acknowledgements

This work was supported by the Novo Nordisk Foundation.

## Author contributions

E.D.J, T.J., J.D.K. and M.K.J. conceived this project. E.D.J, T.J. L.E.P. and M.K.J. designed all of the experiments. E.D.J., M.L., and T.J. performed all growth and evolution assays. E.D.J., and M.L constructed all strains and plasmids. E.D.J, T.J. L.E.P., and M.K.J analyzed the data. E.D.J and M.K.J. wrote the paper.

## Materials and correspondence

Inquiries about materials should be sent to corresponding author Michael Krogh Jensen

## Supporting information

**Supplementary Fig. S1**. Representative plates showing red/white frequencies of yeast colonies with or without T7RNAP-mediated expression of gRNA and cgRNA targeting Cas9 or iCas9 to *ADE2* locus.

**Supplementary Fig. S2**. Frequencies of red colony formations in yeast cells with plasmids encoding three different terminator designs upstream the expression cassette encoding *ADE2*_cgRNA. Frequencies of red yeast colony formation when transforming yeast with each of three different *ADE2*_cgRNA-expressing plasmids together with improved Cas9 (iCas9) in the absence (-) or presence (+) of T7RNAP are shown as mean ± s.d. from three (*n* = 3) biological replicate experiments.

**Supplementary Fig. S3**. T7RNAP^F11L/T613A^ performs better in yeast than wild-type T7RNAP. (**A**) Mean fluorescence intensity (MFI) ± s.d of T7RNAP:GFP fusions and a negative control (empty plasmid) based on 10,000 single cell events from three (*n* = 3) biological replicates. (**B**) Frequency of *ADE2* disruption phenotype when using T7RNAP wild-type and mutant variants for *ADE2*_cgRNA expression. Data show mean frequencies of red colony formation ± s.d. from three (*n* = 3) biological replicate experiments. Significance was determined relative to wild-type T7RNAP from biological triplicates, where **p<0.005, and N.S. = not significant.

**Fig. S4**. dCas9 facilitates cgRNA-DNA editing. Resistant colonies expressing T7RNAP^F11L/T613A^ and (**A**) Cas9 or (**B**) dCas9 with cgRNA where indicated (+CAN1_cgRNA) on plates containing L-Canavanine (600 μg/ml) following 72 hrs of liquid incubation. Parental strain (Sc36 with closed auxotrophies) was plated in parallel for comparison. (**C**) Quantification of resistant colonies per viable cells. Controls without cgRNA expressed were pooled from strains expressing Cas9 or dCas9 and from strain Sc36 with closed auxotrophies. Error bars are shown as mean ± s.d. from three (*n* = 3) biological replicate experiments and significance determined, where * p<0.05 and ** p<0.005. Colony scores are presented in Supplementary Table S1.

**Fig. S5**. Plasmid sequencing shows no sign of mutagenesis in mutant strains. **(A)** Schematic outline of workflow as in Fig. 4. Cas9 and T7RNAP^F11L/T613A^ (epT7RNAP) are expressed during growth in galactose and together target the chromosomal locus for engineering with mutant cgRNA. **(B)** Plasmids from the induction workflow shown in Fig. 4 were Sanger sequenced (n = 3). The reverse orientation of *cgRNA_HIS3_stop* is shown. Chromatograms are presented with blue shading indicating the engineered STOP codon (sense: AAG -> TAG). (**C**) Identified mutations from Fig. 4B were introduced in the *HIS3* cassette of the empty vector pEDJ437 (pEDJ510-513) and transformed into clean strains (CEN.PK2-1C). Wild-type (WT) *HIS3* contained in pEDJ437 (top-left) displays viable colonies on synthetic complete histidine dropout plates, while the engineered *HIS3* STOP codon contained in pEDJ509 is not viable. Mutants contained in pEDJ510-513 all give rise to viable *HIS3* mutants.

**Table S1**. Colony forming units, OD600 measurements, and calculations of mutation frequencies and rate.

**Table S2**. List of strains used in this study.

**Table S3**. List of plasmids used in this study.

**Table S4**. List of primers used in this study.

**Table S5**. List of gRNA sequences used in this study.

**Table S6**. List of gBlocks and assembly parts used in this study.

