## Supplementary tables for "A synthetic RNA-mediated evolution system in yeast"

**Supporting Information**

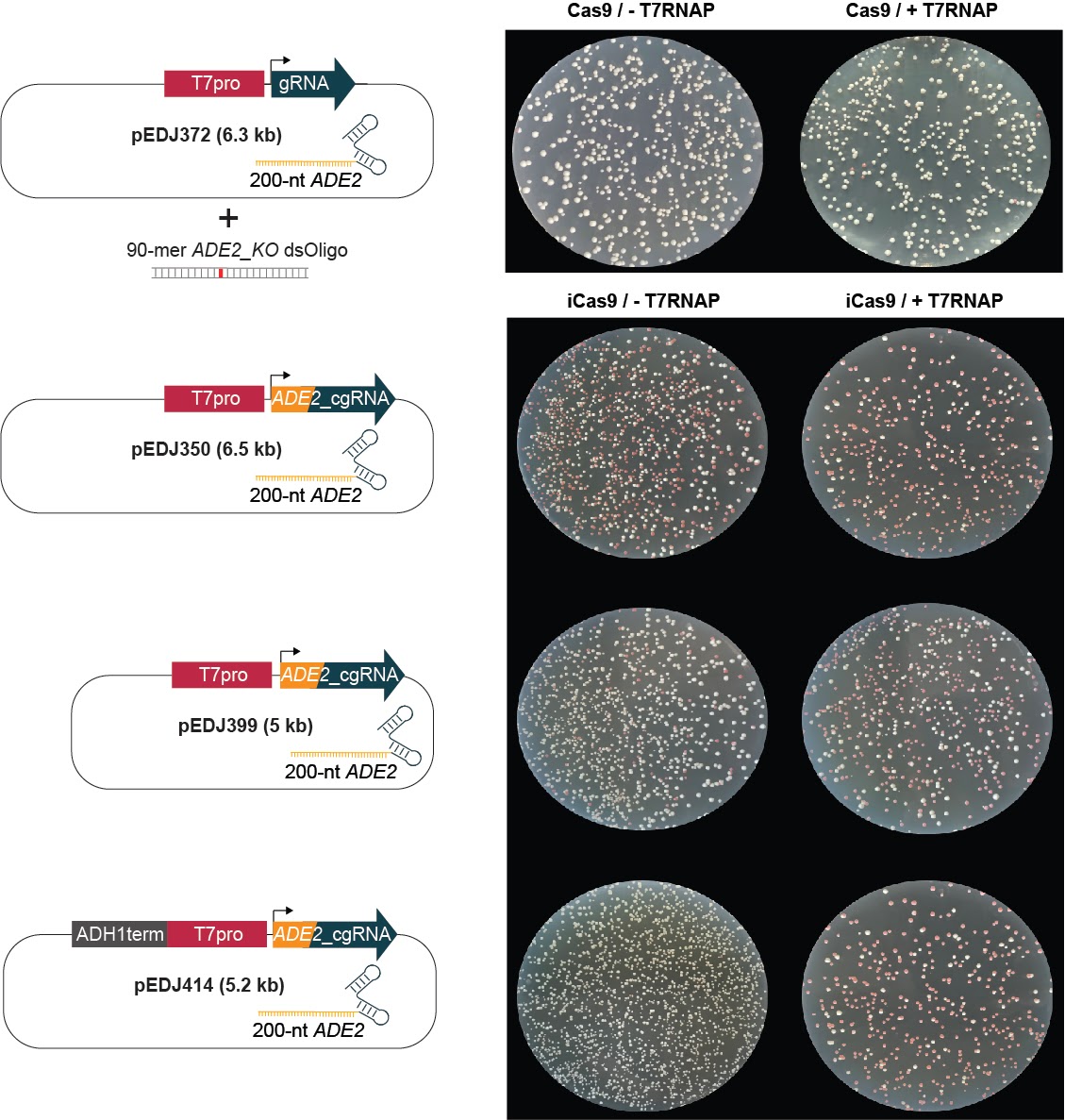

**Supplementary Fig. S1.** Representative plates showing red/white frequencies of yeast colonies with or without T7RNAP-mediated expression of gRNA and cgRNA targeting Cas9 or iCas9 to ADE2 locus

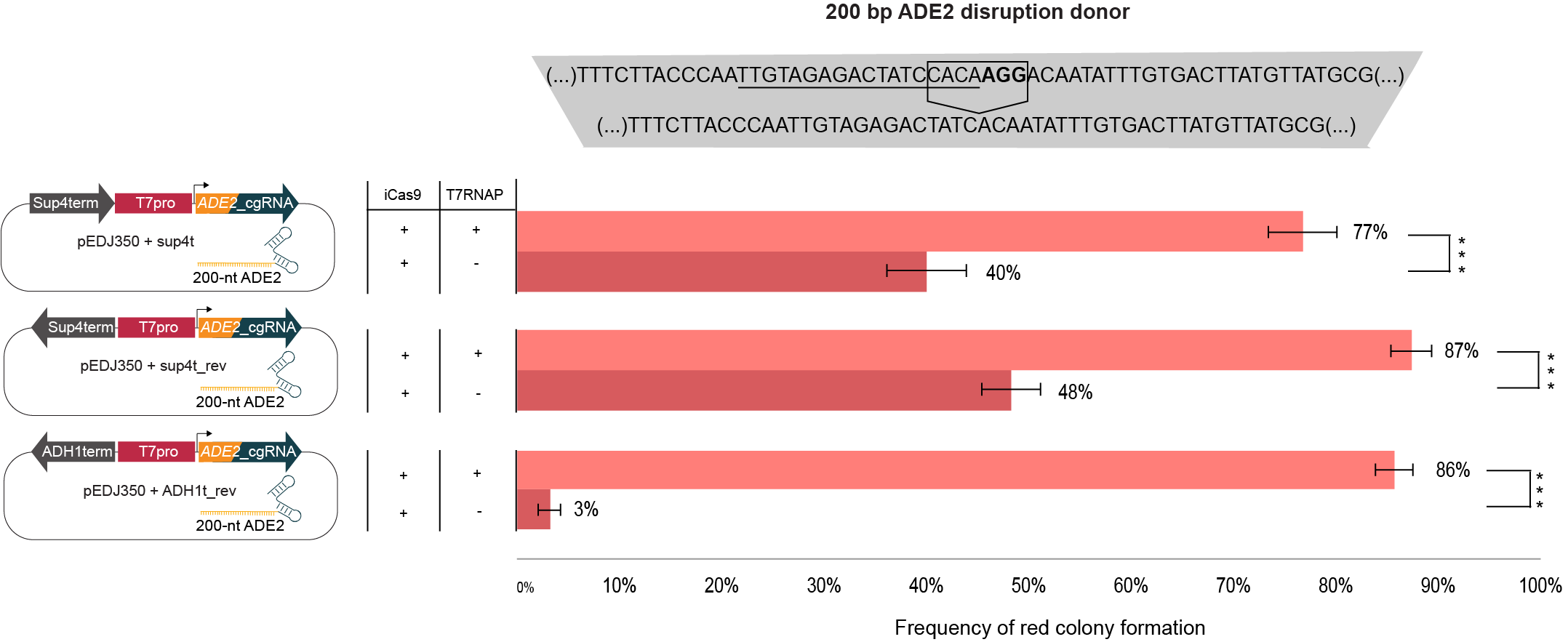

**Supplementary Fig. S2.** Frequencies of red colony formations in yeast cells with plasmids encoding three different terminator designs upstream the expression cassette encoding *ADE2*_cgRNA. Frequencies of red yeast colony formation when transforming yeast with each of three different *ADE2*_cgRNA-expressing plasmids together with improved Cas9 (iCas9) in the absence (-) or presence (+) of T7RNAP are shown as mean ± s.d. from three (*n* = 3) biological replicate experiments.

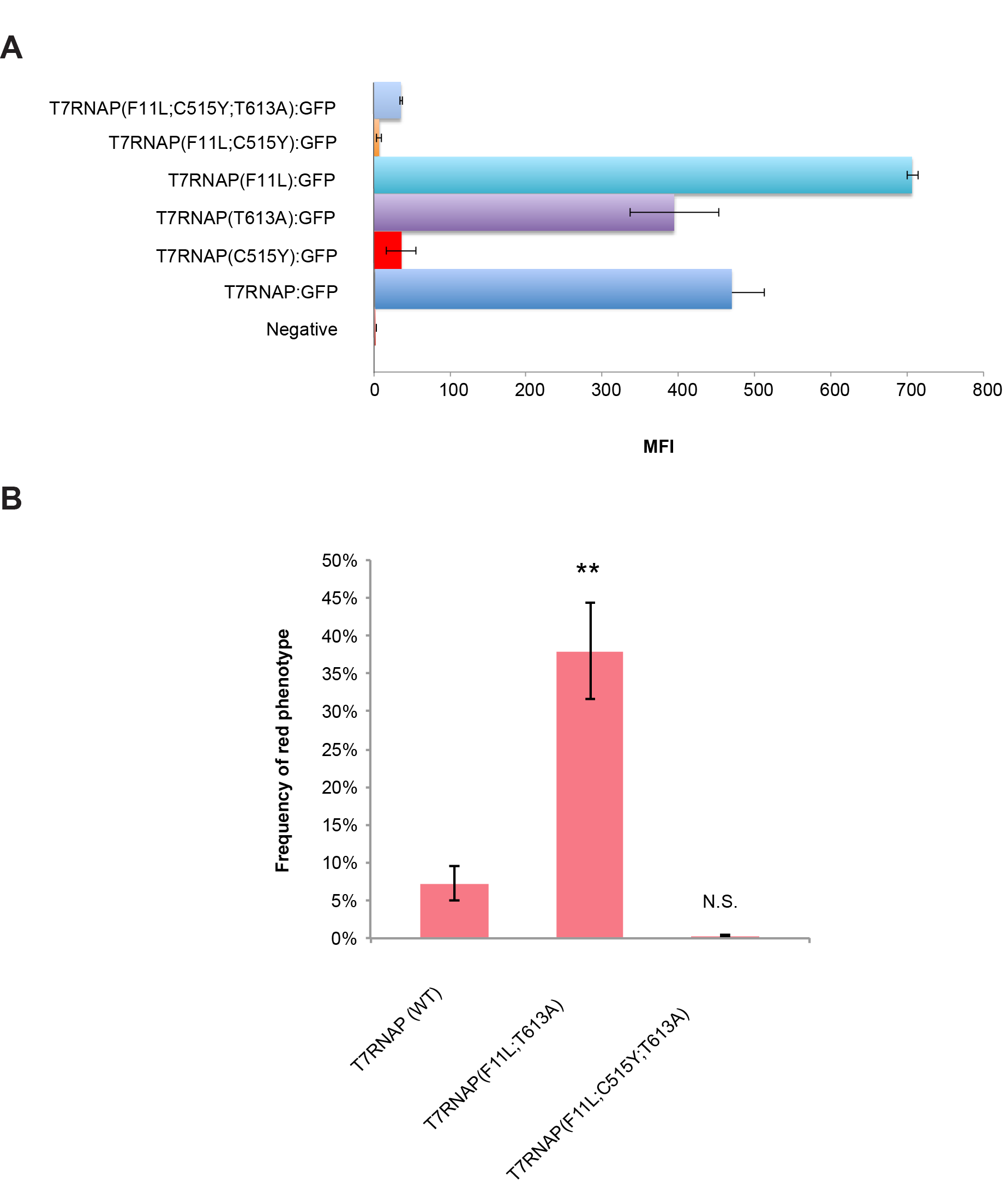

**Supplementary Fig. S3. T7RNAP^F11L/T613A^ performs better in yeast than wild-type T7RNAP.** (**A**) Mean fluorescence intensity (MFI) ± s.d of T7RNAP:GFP fusions and a negative control (empty plasmid) based on 10,000 single cell events from three (*n* = 3) biological replicates. (**B**) Frequency of *ADE2* disruption phenotype when using T7RNAP wild-type and mutant variants for *ADE2*_cgRNA expression. Data show mean frequencies of red colony formation ± s.d. from three (*n* = 3) biological replicate experiments. Significance was determined relative to wild-type T7RNAP from biological triplicates, where **p<0.005, and N.S. = not significant.

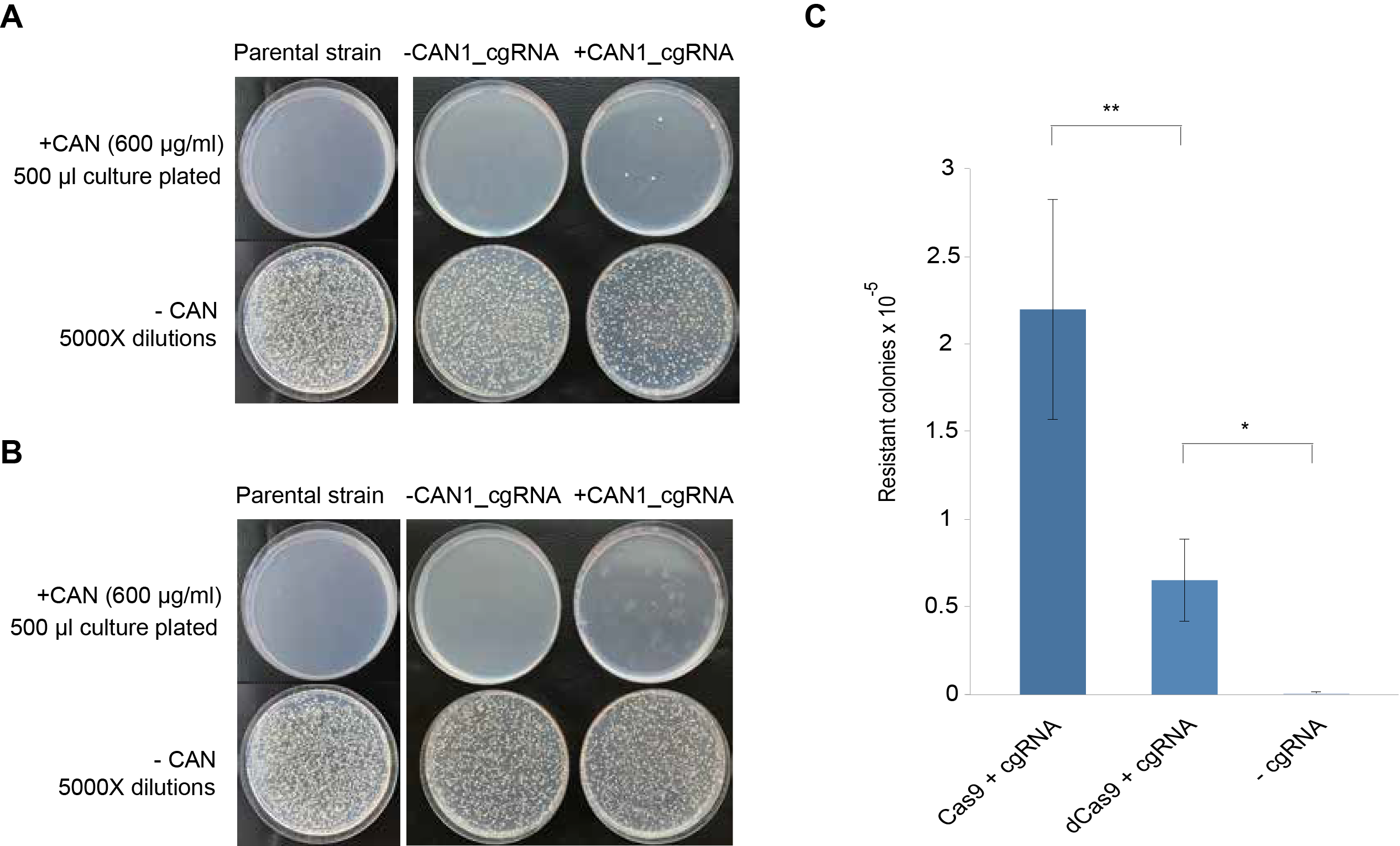

**Fig. S4. dCas9 facilitates cgRNA-DNA editing**. Resistant colonies expressing T7RNAP^F11L/T613A^ and (**A**) Cas9 or (**B**) dCas9 with cgRNA where indicated (+CAN1_cgRNA) on plates containing L-Canavanine (600 μg/ml) following 72 hrs of liquid incubation. Parental strain (Sc36 with closed auxotrophies) was plated in parallel for comparison. (**C**) Quantification of resistant colonies per viable cells. Controls without cgRNA expressed were pooled from strains expressing Cas9 or dCas9 and from strain Sc36 with closed auxotrophies. Error bars are shown as mean ± s.d. from three (*n* = 3) biological replicate experiments and significance determined, where ** p<0.05 and *** p<0.005. Colony scores are presented in Supplementary Table S1.

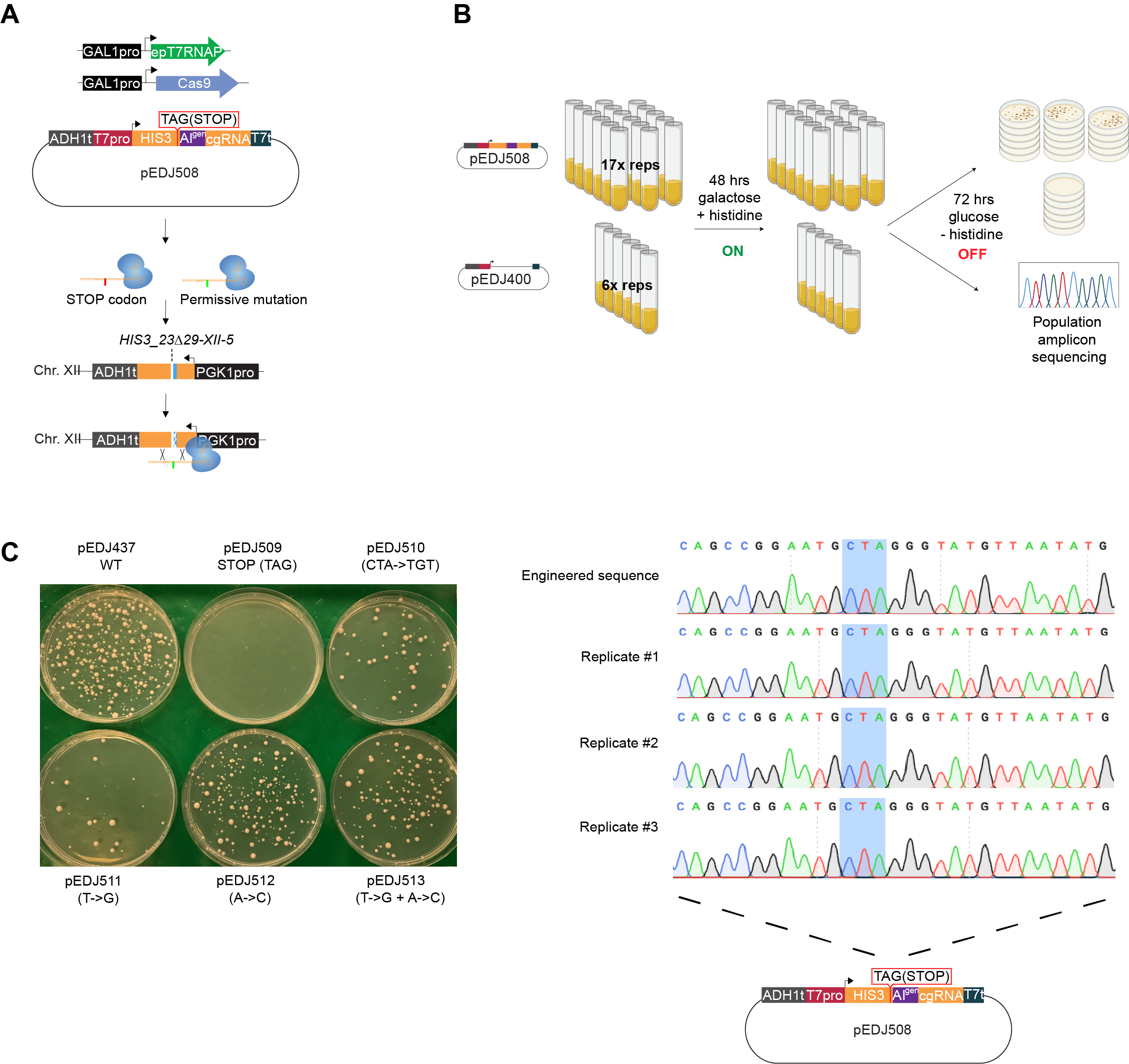

**Fig. S5. Plasmid sequencing shows no sign of mutagenesis in mutant strains. (A)** Schematic outline of workflow as in Fig. 4. Cas9 and T7RNAP^F11L/T613A^ (epT7RNAP) are expressed during growth in galactose and together target the chromosomal locus for engineering with mutant cgRNA. **(B)** Plasmids from the system induction workflow shown in Fig. 4 were Sanger sequenced. The reverse orientation of *cgRNA_HIS3_stop* is shown. Chromatograms are presented with blue shading indicating the engineered STOP codon (sense: AAG -> TAG). (**C**) Identified mutations from Fig. 4B were introduced in the *HIS3* cassette of the empty vector pEDJ437 (pEDJ510-513) and transformed into clean strains (CEN.PK2-1C). Wild-type (WT) *HIS3* contained in pEDJ437 (top-left) displays viable colonies on synthetic complete histidine dropout plates, while the engineered *HIS3* STOP codon contained in pEDJ509 is not viable. Mutants contained in pEDJ509-513 all give rise to viable *HIS3* mutants.
